# Plasma Metabolomic Profiling of COPD Patients Stratified by Smoking Status: A GC-MS-Based Approach

**DOI:** 10.64898/2026.08.12.744361

**Authors:** Rishita Singh, Sudip Ghosh, Amit Kumar Mandal

## Abstract

**Background:** Chronic obstructive pulmonary disease, primarily caused by exposure to cigarette smoke, is a heterogeneous lung condition characterized by complex metabolic alterations. The metabolic changes associated with smoking status have not been thoroughly investigated. Our study aims to explore the metabolite profile of COPD patients categorised by their smoking habits, including smokers, ex-smokers, and non-smokers.

**Methods:** In this study, the plasma metabolome of smoking stratified COPD patients were assessed using gas chromatography coupled to mass spectrometry. We applied multivariate and univariate statistical analysis to identify the differentially abundant metabolites.

**Results:** We identified 23 altered metabolites in the smokers and 36 in the ex-smokers COPD subgroups. Interestingly, in comparison to the control group, no significant alteration was observed in the plasma of non-smoker COPD patients. Additionally, pathway enrichment analysis revealed top dysregulated metabolic pathways, including biosynthesis of unsaturated fatty acids, galactose metabolism, phenylalanine, tyrosine, and tryptophan biosynthesis, and glycosylphosphatidylinositol (GPI)-anchor biosynthesis. The receiver operating characteristic curve screened five metabolites, such as tetradecanoic acid, 2,4-di-tert-butylphenol, chloroxylenol, tetradecanal, and 1-dodecene, with the highest diagnostic performance (AUC > 0.8).

**Conclusion:** This study reveals distinct plasma metabolic signatures across COPD subgroups categorized by cigarette smoking history.

## 1. Introduction

Chronic obstructive pulmonary disease (COPD) is a leading cause of morbidity and mortality worldwide, imposing a substantial public health burden, particularly in low- and middle-income countries. COPD is a heterogeneous respiratory disorder characterized by persistent airflow limitation resulting from structural and functional abnormalities in the airways and lung parenchyma, including chronic bronchitis, small airway disease (bronchiolitis), emphysema, and pulmonary vascular remodeling. These pathological changes are associated with symptoms such as chronic cough, sputum production, wheezing, and progressive dyspnea, ultimately impairing expiratory airflow. The resulting airflow obstruction is typically assessed by spirometry, which demonstrates a reduced forced expiratory volume in one second (FEV₁) and a decreased FEV₁/forced vital capacity (FVC) ratio. Although spirometry remains the gold standard for the diagnosis of COPD, it has several limitations. These include limited sensitivity for detecting early small airway dysfunction, an inability to distinguish between different COPD phenotypes, the potential for misclassification due to the inherent limitations of lung function measurements, and the risk of overdiagnosis in older individuals when the fixed FEV₁/FVC ratio (<0.70) is used for diagnosis **[1,2,3].** Consequently, these challenges have prompted increasing interest in the identification of novel molecular markers for the early diagnosis and precise detection of COPD. There are many causal factors of COPD, including environmental factors such as tobacco smoke, exposure to biomass/fossil fuel burning, and genetic factors, i.e, α1-antitrypsin deficiency. Cigarette smoking (CS) is one of the major risk factors for COPD development and progression, causing approximately 90% of total COPD cases **[4]**. CS is a mixture of approximately 4500 noxious chemical particles, which, upon entering the human body, activate a cascading immune response and generation of reactive oxidants, resulting in impairment of downstream physiological and biochemical processes **[5,6]**. The role of CS in COPD development and progression is well established, yet the molecular signatures related to CS in COPD individuals stratified by smoking status are unknown to date. Omics-based studies provide a platform to uncover molecular signatures associated with diseases. Previous omics studies in COPD have highlighted widespread metabolic dyregulation in plasma, sputum, saliva, and urine samples. These studies have reported alterations in energy metabolism, arginine metabolism, lipid remodeling, and redox imbalance, etc **[1,7,8,9,10,11,12]**. Plasma, being a complex biofluid, has been widely used to explore many systemic diseases and their associated biomarkers **[13,14]**. However, the knowledge gap persists regarding CS-associated metabolic alterations in the plasma sample of COPD patients. Therefore, elucidating the molecular signature of COPD patients with varying degrees of cigarette smoking is crucial for advancing early and precise interventions. In our previous study, we reported metabolic alterations in the saliva of smokers, ex-smokers, and non-smokers categories of COPD patients. However, plasma as a central and integrating matrix has the potential to reveal detailed metabolic perturbations in COPD.

Our study attempts to fill this gap by exploring the plasma metabolome of COPD patients categorised by smoking status, such as smokers, ex-smokers, and non-smokers. We aimed to elucidate the metabolic alterations in COPD patients with different smoking habits and to investigate metabolic markers associated with COPD to better understand disease pathology and molecular phenotyping.

## 2. Materials and methods

### 2.1. Materials

All chemicals of analytical grade, such as Methoxyamine hydrochloride (MOX), N-Methyl-N-(trimethylsilyl)trifluoroacetamide (MSTFA), Trimethylchlorosilane (TMCS), and Ribitol, were purchased from Sigma-Aldrich (USA). LC-MS grade methanol, water, and HPLC-grade hexane and pyridine were purchased from Thermo Fisher Scientific (USA).

### 2.2. Study design

The study was conducted at the Indian Institute of Science Education and Research (IISER), Kolkata, in collaboration with the All India Institute of Medical Sciences (AIIMS), Kalyani. The study was approved by the institutional ethics committee, with reference no. IEC/AIIMS/Kalyani/certificate/2024/415. Participants with a known case of COPD were recruited into the study after obtaining their consent. Patients with and without the habit of smoking were considered for the study after a full inquiry with them. COPD is diagnosed after a preliminary Spirometry test in the hospital setting (AIIMS-Kalyani), and the criterion for inclusion was post FEV_1_/FVC < 70%. Following spirometry, blood samples were collected from COPD patients (n=30) and healthy controls (n=8). These COPD patient samples were further categorised into Smoker (n=15), Ex-smoker (n=12), and non-smoker (n=3).

### 2.3. Sample collection

2 mL venous blood sample was collected in an EDTA-coated vacutainer tube. The blood sample was centrifuged at 3000 rpm for 10 min at 4 °C to separate plasma. Plasma samples obtained from the supernatant were then aliquoted and stored at -80 °C until analysis. The stored plasma samples were thawed at 4 °C, and 100 μL of the sample was taken for metabolite extraction using methanol and water. The collected supernatant was lyophilised and derivatized using the protocol described elsewhere **[9]**.

### 2.4. GC-MS Analysis

The plasma metabolites were analysed on an Agilent 8890 GC system coupled with a 7010C GC/Triple Quad MS system (Agilent Technologies, USA). The separation of metabolites was conducted using a DB-5MS column, coated with 5% phenyl-methylpolysiloxane (30 m x 250 μm i.d., x 0.25 μm), Agilent Technologies, USA. The initial temperature of the GC oven was set at 60 °C for 1 minute, followed by an increase at a rate of 10 °C per minute until reaching 325 °C. The temperatures for the inlet, transfer line, and ion source were maintained at 250 °C, 290 °C, and 250 °C, respectively. A sample injection volume of 1 μL was used with a split ratio of 1:5, and helium served as the carrier gas at a flow rate of 1 mL per minute. Data were acquired in full scan mode over a range of 50-800 m/z, with ions generated using electron impact ionization at 70 eV.

### 2.5. Data Processing and Statistical Analysis

Agilent MassHunter workstation software (Agilent Technologies, USA) was used to process the raw data. The metabolite identities were annotated based on a spectral match score ≥60 against the National Institute of Standards and Technology (NIST) library. The metabolites with >50% missing values per group were removed, and imputation was performed with one-fifth of the minimum positive value. The data was transformed and scaled using MetaboAnalyst 6.0 software. Further, Principal Component Analysis (PCA), Partial Least Squares Discriminant Analysis (PLSDA), Orthogonal Partial Least Squares Discriminant Analysis (O-PLSDA), and univariate Wilcoxon rank-sum test with Benjamini-Hochberg correction, was performed to identify significant features based on variable importance in projection (VIP>1), fold change (FC ≥2), and FDR-adjusted p-value <0.05. The altered metabolites with VIP score >1, p-value <0.05, and FC ≥2 were taken for pathway enrichment and disease enrichment analysis using Kyoto Encyclopedia of Genes and Genomes (KEGG) and the Human Metabolome Database (HMDB).

The metabolites obtained from multivariate and univariate analyses were evaluated for their diagnostic potential using random forest analysis with an AUC > 0.8. The metabolite-gene-disease network was constructed using Cytoscape 3.10.3 to investigate the relationships between identified differential metabolites and COPD-associated genes obtained from the Comparative Toxicogenomics Database (CTD). Furthermore, Spearman’s rank correlation analysis was performed to evaluate the associations between metabolite levels and spirometric indices.

## 3. Results

### 3.1. General Characteristics of the Participants

The study encompasses 30 patients diagnosed with COPD, categorized into the following subgroups: 15 COPD smokers, 12 COPD ex-smokers, 3 COPD non-smokers, and 8 healthy controls. The demographic details and clinical parameters were analyzed and are presented as mean values with standard deviations. Statistical parameters were derived using GraphPad Prism 8.0.2, as shown in **Table 1**.

**Table 1.** Clinical parameters of COPD patients and healthy controls.

| Patient's clinical details | COPD patient<br>Smoker<br>(n=15) | COPD patient<br>Ex-smoker<br>(n=12) | COPD patient<br>Non-smoker<br>(n=3) | Healthy<br>Non-smoker<br>(n=8) |
| --- | --- | --- | --- | --- |
| **Age** | 60.60 ± 8.41 | 61.85 ± 8.99 | 49.67 ± 17.2 | 25.63 ± 3.15 |
| **Sex (M/F)** | 15/0 | 12/0 | 1/2 | 6/2 |
| **Height (cm)** | 159.4 ± 6.15 | 162.3 ± 4.94 | 153.7 ± 2.0 | 166.3 ± 6.45 |
| **Weight (Kg)** | 54.0 ± 10.73 | 61.45 ± 10.87 | 48.0 ± 1.7 | 67.18 ± 14.9 |
| **BMI** | 21.22 ± 3.65 | 23.45 ± 4.56 | 20.3 ± 1.2 | 24.7 ± 4.7 |
| **Cigarette (Pack years)** | 42.95 ± 23.87 | 25.3 ± 27.46 | N/A | N/A |
| **Smoking years** | 34.07 ± 8.46 | 24.17 ± 13.05 | N/A | N/A |
| **Pre-FEV<sub>1</sub>** | 0.93 ± 0.40 | 0.81 ± 0.32 | 1.13 ± 0.30 | 3.06 ± 0.73 |
| **Post- FEV<sub>1</sub>** | 1.11 ± 0.41 | 0.86 ± 0.32 | 1.20 ± 0.32 | 3.18 ± 0.75 |
| **Pre- FVC** | 2.06 ± 0.63 | 1.9 ± 0.67 | 2.42 ± 0.66 | 3.59 ± 0.96 |
| **Post- FVC** | 2.26 ± 0.69 | 2.04 ± 0.74 | 2.34 ± 0.69 | 3.6 ± 0.95 |
| **Pre- FEV<sub>1</sub>/FVC** | 44.88 ± 10.8 | 45.3 ± 18.3 | 46.80 ± 1.35 | 85.58 ± 5.4 |
| **Post- FEV<sub>1</sub>/FVC** | 48.6 ± 15.46 | 44.7 ± 16.9 | 51.53 ± 4.31 | 88.3 ± 5.0 |
| **All data points are Mean ± SD, n = number of samples, FEV<sub>1</sub> = Forced expiratory volume in 1 sec, FVC = Forced vital capacity** |  |  |  |  |

### 3.2. Comparative metabolic Profiling between different COPD subgroups

The multivariate PCA and PLS-DA model separated the COPD subgroups (smokers, ex-smokers, and nonsmokers) from controls **(Fig 1A, B)**. Upon cross-validation, the model shows a goodness of fit (R² = 0.80) and predictive capability (Q² = 0.67), with a permutation (n=2000) p-value of 0.0015, indicating strong separation between the groups. The hierarchical clustering heatmap and variable-of-importance projection obtained from PLS-DA show the features that segregate the groups **(Fig. 2A, B)**. The Pairwise comparison of smoker vs control **(Fig 1C, D)**, ex-smoker vs control **(Fig 1E, F)**, and non-smoker vs control **(Fig 1G, H)** represents R²Y = 0.95 and Q² = 0.90 (p = 5e-04), R²Y = 0.96 and Q² = 0.93 (p = 5e-04), R²Y = 0.63 and Q² = 0.16 (p = 0.2) respectively. The univariate Kruskal-Wallis with Benjamini-Hochberg correction of COPD subgroups vs controls identifies 18 metabolites with P_Adj_ <0.05, **Table S1**. Further, for the pairwise comparison, univariate Wilcoxon rank-sum test with the Benjamini-Hochberg correction identified 23 differential metabolites in the smokers subgroup **Table S2**, 36 in the ex-smokers subgroup **Table S3**, with P_Adj_ <0.05, VIP > 1, FC ≥2, AUC > 0.8, whereas no significant metabolites were observed in the non-smokers subgroup. The volcano plot and VIP score plot in the smokers **(Fig 2C, D)**, ex-smokers **(Fig 2E, F)**, represent differentially expressed metabolites, whereas non-smokers show no significant differential expression **(Fig 2E, F)**.

**Figure 1.**
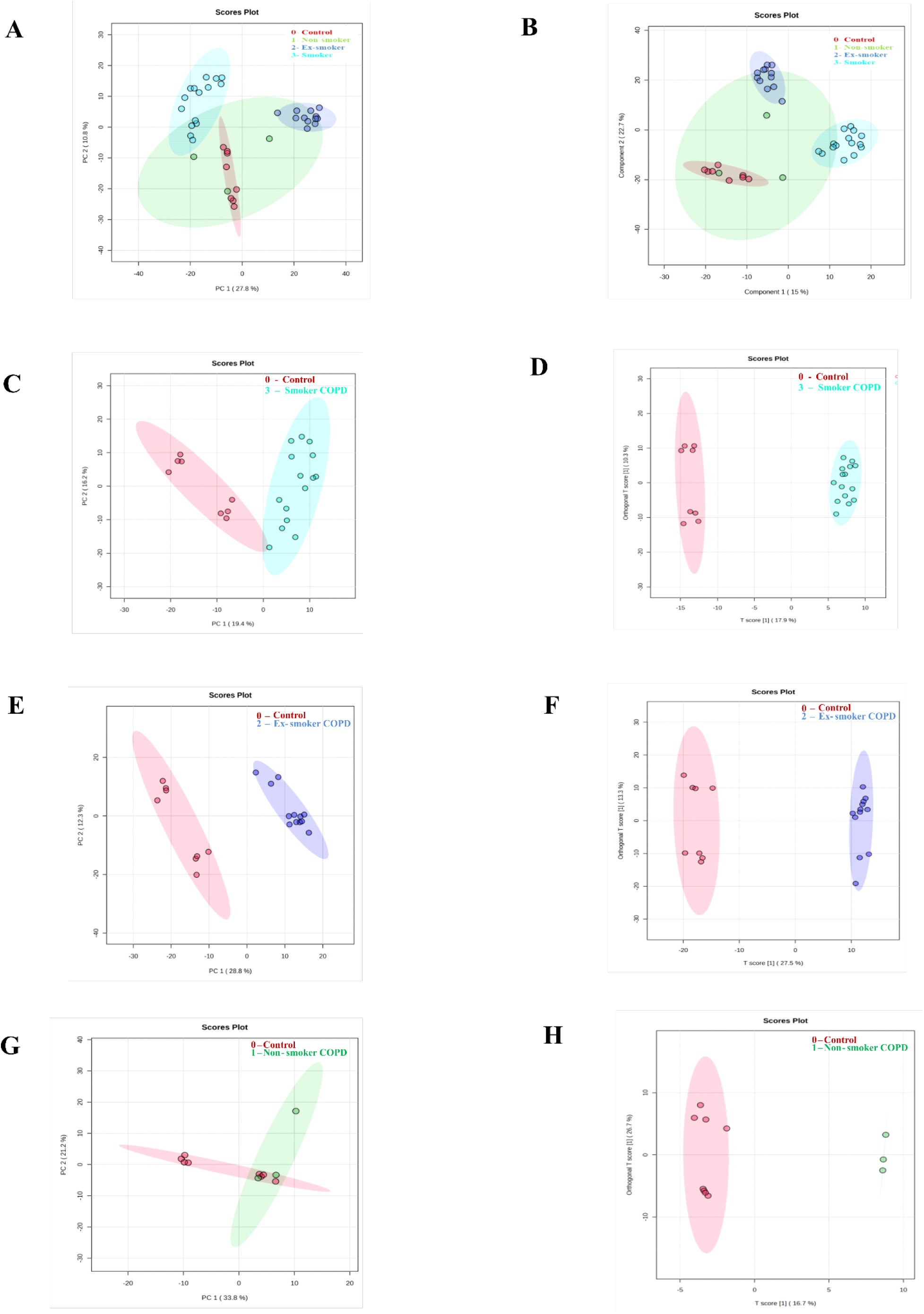
Multivariate analysis of plasma metabolites across COPD subgroups, (A) PCA score plot between COPD subgroups and control (green: Non-smoker COPD, blue: Ex-smoker COPD, cyan: Smoker COPD, red: Healthy control), (B) PLSDA score plot of the respective subgroups, (C) PCA plot of smokers vs control (D) OPLSDA plot of smokers vs control, (E) PCA analysis between ex-smokers vs control, (F) OPLSDA between ex-smokers vs control, (G) PCA plot of non-smokers vs control, (H) OPLSDA between non-smokers vs controls.

**Figure 2.**
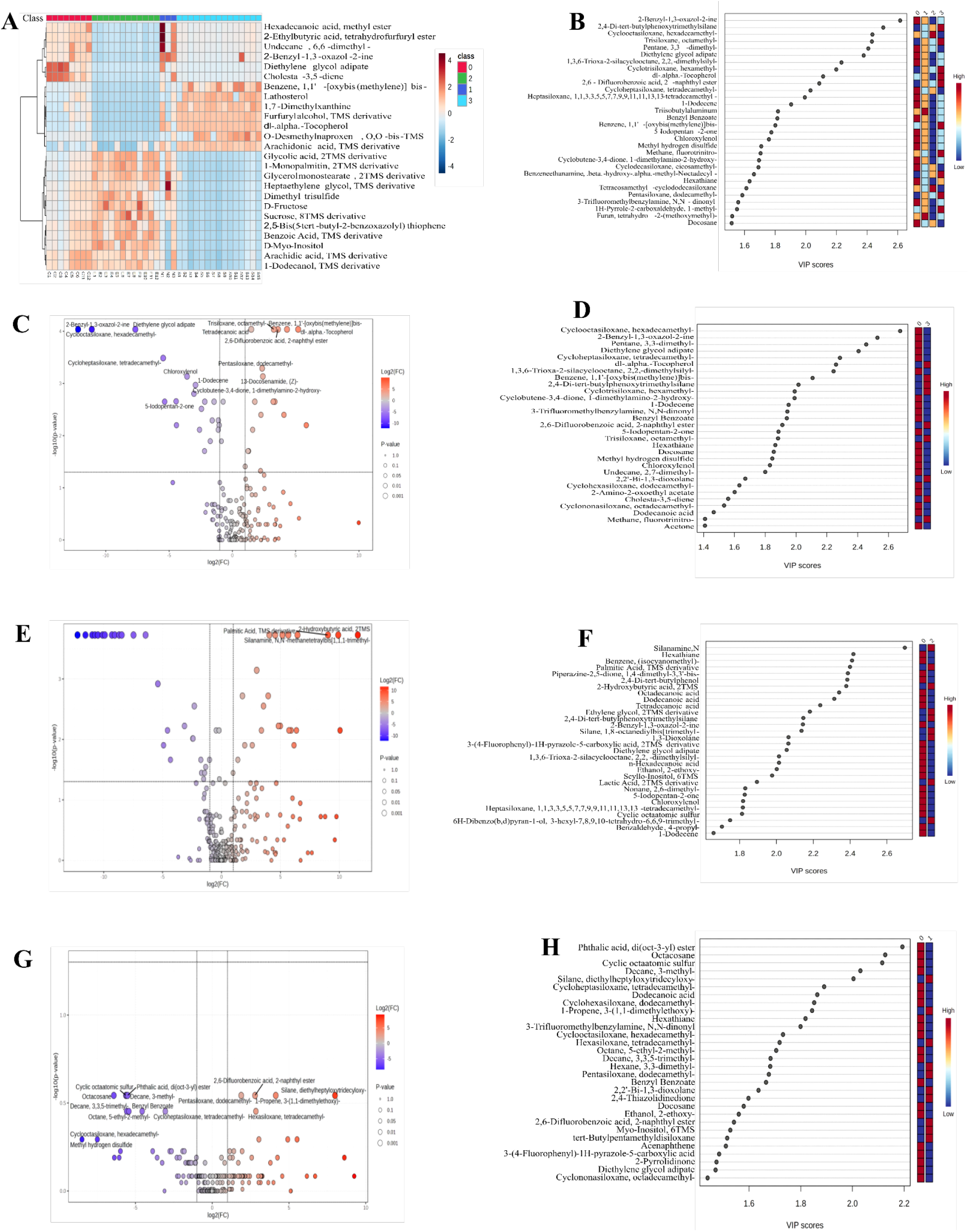
Visualisation of variable of importance from volcano and VIP plot (A) Heatmap depicting hierarchical clustering of metabolites in different subgroups (0:control, 1:non-smokers, 2:ex-smokers, 3:smokers) (B) VIP score plot obtained from PLSDA analysis in COPD vs control, (C) Volcano plot of differential metabolites with -log_10_(p-value) and Log_2_FC in smokers, (D) VIP score plot obtained from OPLSDA in smokers subgroup, (E) Volcano plot of differentially expressed metabolites in ex-smokers, (F) VIP plot of ex-smokers subgroup, (G) Volcano plot with no differential expression in non-smokers subgroup, (H) VIP score plot of the non-smokers.

### 3.3. Pathway and Disease Enrichment Analysis

Upon performing enrichment analysis of COPD subgroups using KEGG and HMDB databases, we revealed disrupted pathways such as biosynthesis of unsaturated fatty acids, galactose metabolism, phenylalanine, tyrosine, and tryptophan biosynthesis, glycosylphosphatidylinositol (GPI)-anchor biosynthesis, etc **(Fig 3A)**. COPD smoker subgroup highlighted perturbation in fatty acid biosynthesis, glycosylphosphatidylinositol (GPI)-anchor biosynthesis, and biosynthesis of unsaturated fatty acids **(Fig 3C)**. In contrast, COPD ex-smokers show alterations in fatty acid biosynthesis, pyruvate metabolism, glycolysis/gluconeogenesis, and biosynthesis of unsaturated fatty acids **(Fig 3E)**. Additionally, the disease enrichment analysis identifies overrepresented diseases such as schizophrenia, colorectal and pancreatic cancer, epilepsy, etc., in COPD subgroups **(Fig 3B, 3D, 3F).**

**Figure 3.**
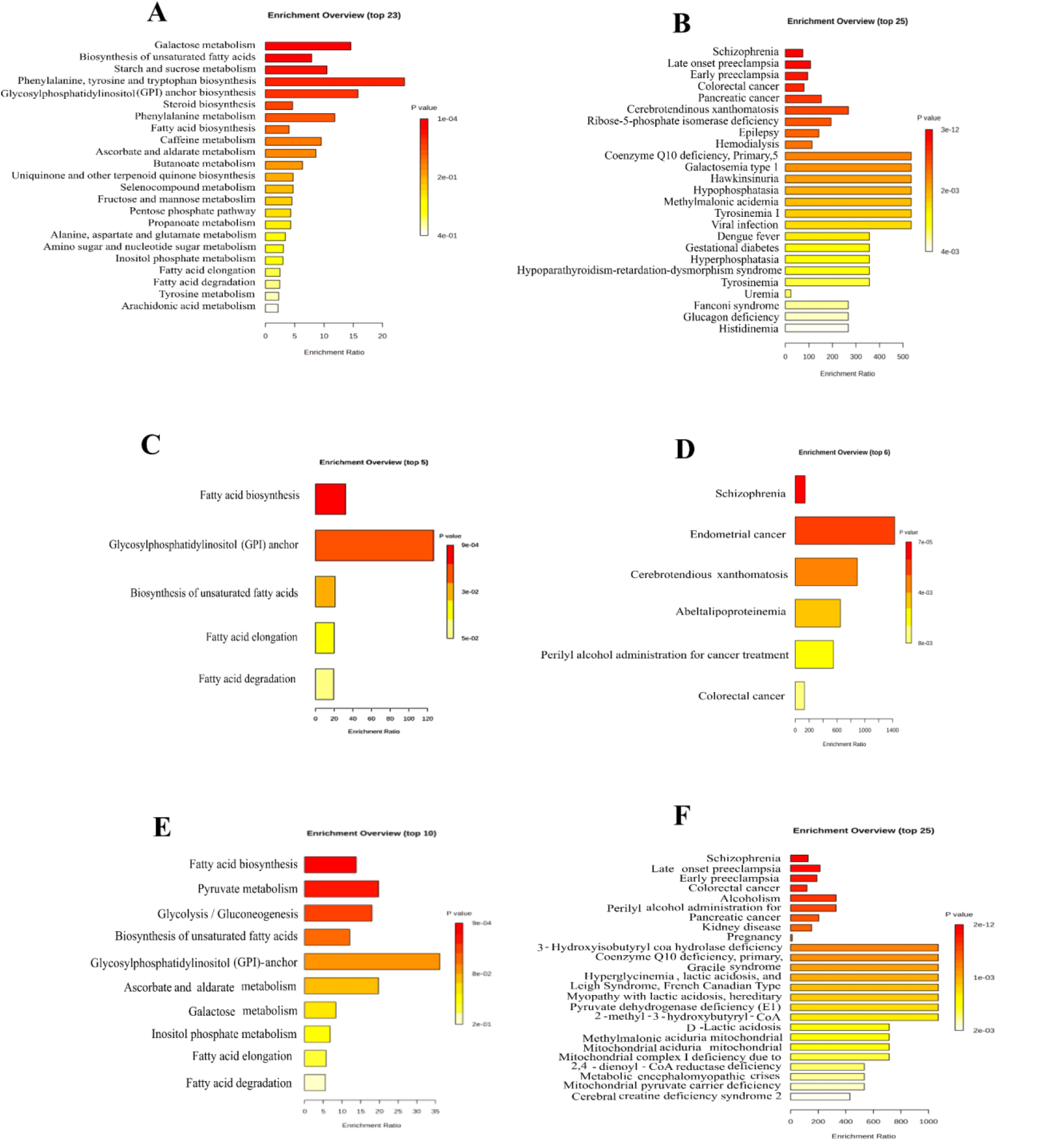
Pathway enrichment and disease enrichment analysis using KEGG and HMDB (A) Perturbed metabolic pathways in COPD, (B) Disease linked with altered metabolites in COPD, (C) Altered pathways in smokers subgroup, (D) Disorders associated with altered metabolome profile in smokers, (E) Perturbed pathways in ex-smokers, (F) Disease enrichment overview in ex-smokers.

### 3.4. Biomarker analysis

The metabolites with differential expression were further evaluated for diagnostic performance using a univariate classical ROC curve. There were 5 metabolites across COPD subgroups with an AUC >0.8 (95%CI) and p < 0.05, such as tetradecanoic acid, 1-dodecene, chloroxylenol, tetradecanal, and 2,4-di-tert-butylphenol **(Figure 4)**. Furthermore, subgroup-specific metabolites with an AUC >0.8 (95%CI) were observed, including benzene,1,1’-[oxybis(methylene)]bis-, alpha tocopherol, 13-Docosenamide, acetophenone, docosane in COPD smokers subgroup **(Figure S3)**. Specific metabolites for ex-smokers with an AUC >0.8 (95% CI) included ethylene glycol, lactic acid, 2-hydroxybutyric acid, palmitic acid, and scylloinositol **(Figure S4)**. The scatter dot plot of these metabolites is given in **Figure 5**.

**Figure 4.**
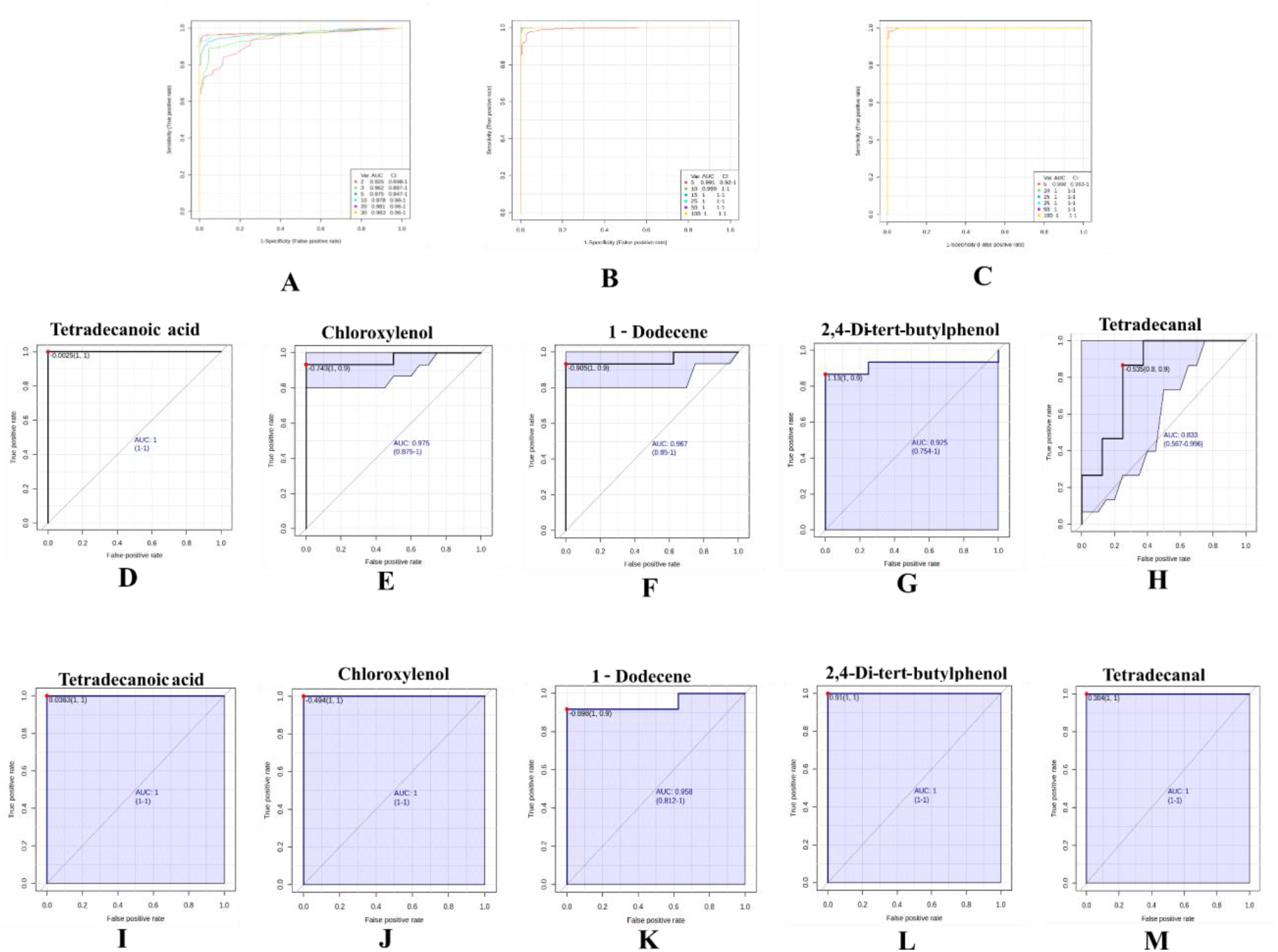
Biomarker analysis of altered metabolites in COPD subgroups, (A-C) Multivariate ROC curve analysis in COPD subgroups, COPD smokers and ex-smokers, respectively, (D-H) Univariate ROC curve of five metabolites with AUC>0.8 in COPD smokers, (I-M) Univariate ROC curve of five metabolites with AUC>0.8 in COPD ex-smokers.

**Figure 5.**
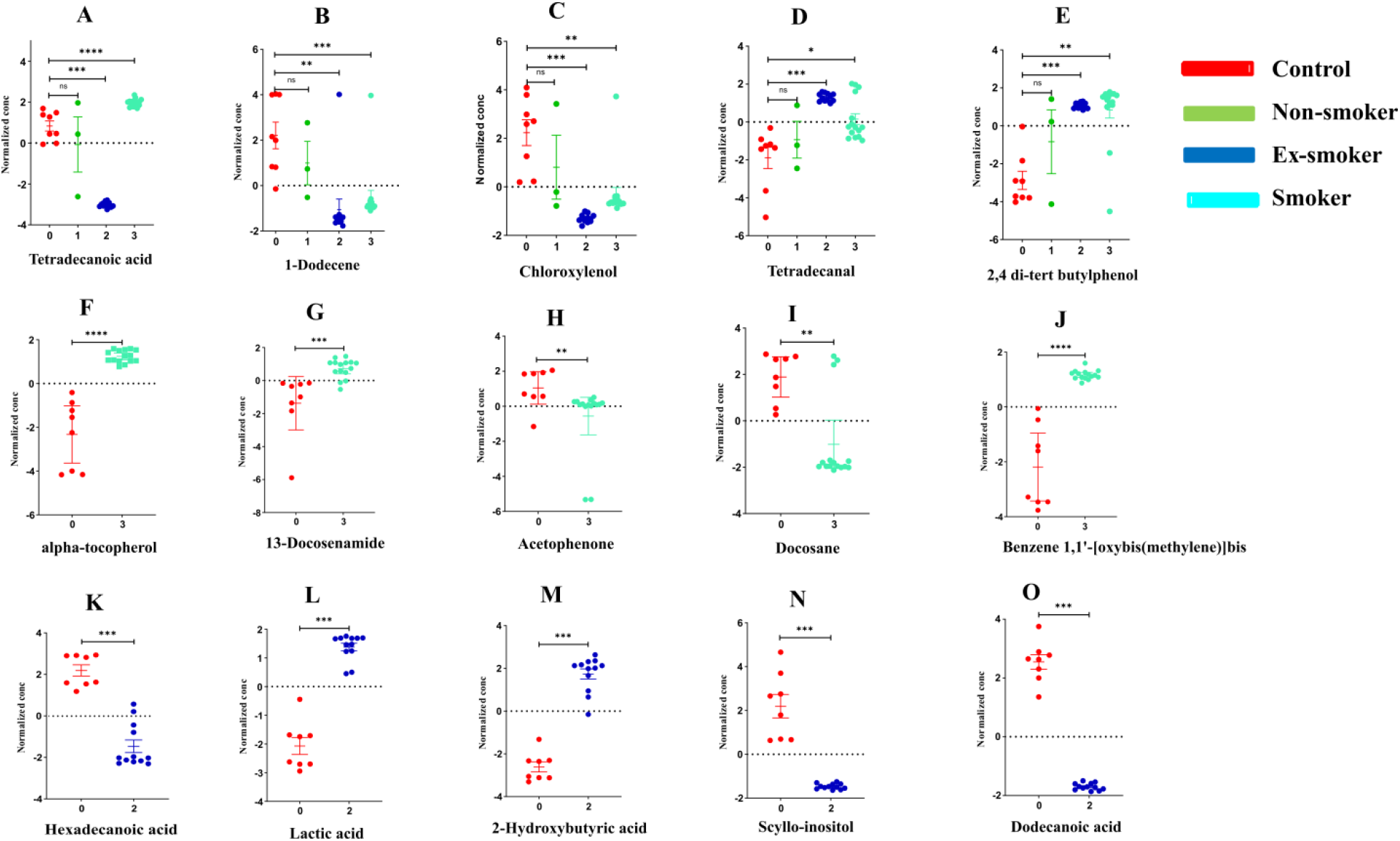
Scatter dot plot of differentially expressed metabolites with high AUC value across different comparisons, (A-E) dysregulated metabolite in both smokers and ex-smokers (red: control, green: non-smokers, blue: ex-smokers, cyan: smokers), (F-J) dysregulated specific to smokers subgroup, (K-O) altered metabolites specific to ex-smokers subgroup, the p-value expressed in the plot is derived from wilcoxon sum rank test with Benjamini-Hochberg correction with *: p<0.05, **: p<0.01, ***: p<0.001, ****: p<0.0001.

### 3.5. Correlation and Gene-metabolite Network Analysis

The Spearman’s rank correlation of differential metabolites is visualised as a heatmap in **Figure 6A**, where 1-dodecene shows a strong positive correlation with chloroxylenol (ρ = 0.88, p < 0.05), and post FEV_1_/FVC (ρ = 0.62, p < 0.05); tetradecanal shows a negative correlation with dodecanoic acid (ρ = -0.5, p < 0.05). In smokers, 1-dodecene shows a positive correlation with chloroxylenol (ρ = 0.88, p < 0.05), post FEV_1_/FVC (ρ = 0.67, p < 0.05), and a negative correlation with tetradecanoic acid (ρ = -0.5, p < 0.05). Tetradecanal showed a positive correlation with 2,4-di-tert-butylphenol (ρ = 0.52, p < 0.05) **(Figure 6B)**. In the ex-smokers subgroup **(Figure 6C),** 1-dodecene exhibited a strong positive correlation with chloroxylenol (p < 0.05, ρ = 0.86), post FEV_1_/FVC (ρ = 0.64, p < 0.05). Moreover, tetradecanal presented a strong positive correlation with 2,4-di-tert-butylphenol ( ρ= 0.91, p < 0.05). Additionally, upon performing the network analysis **(Figure 6D)** we observed several metabolites with a higher degree of centrality, such as alanine, arachidonic acid, palmitic acid, phosphoric acid, stearic acid, tetradecanoic acid, arachidic acid, and α-tocopherol, exhibiting multiple connections with COPD associated genes such as FASN, PLA2G5, PLA2G10, AGXT2, and LGALS13, indicating their potential involvement in key disease related pathways.

**Figure 6.**
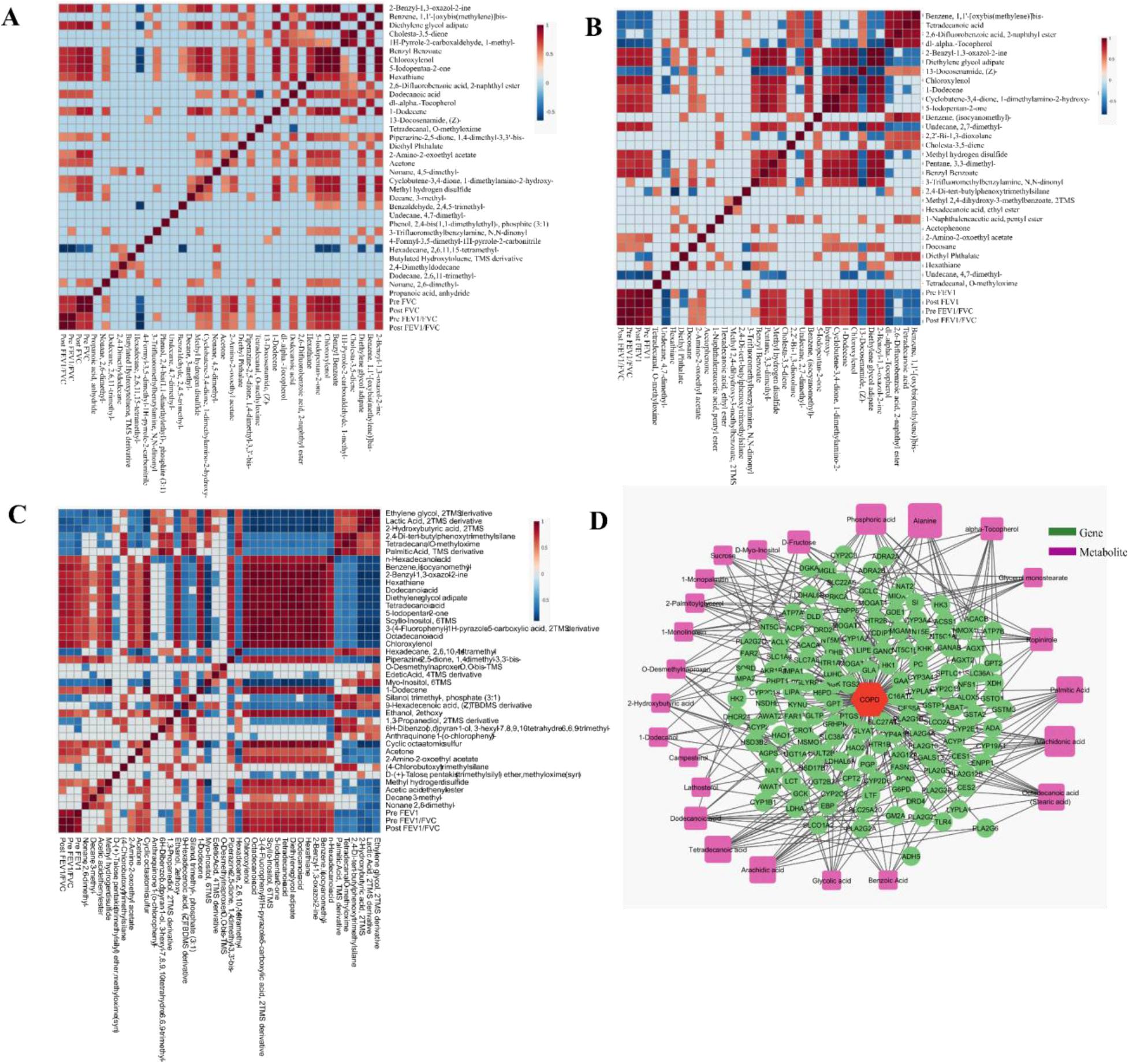
Heatmap representing metabolite and spirometry indices correlation (A) Spearman rank correlation between metabolites and spirometric indices in COPD patients, (B) correlation in COPD smokers (C) correlation in COPD ex-smokers with ρ > 0.5 (D) Metabolite-gene-disease network constructed using differential metabolites identified in COPD patients.

## 4. Discussion

COPD is a heterogeneous disorder usually caused by CS, and it displays many abnormal metabolic processes. However, no study has compared the metabolomic profiles of COPD patients stratified by CS habit. In this study, we investigated the metabolomic profile of smokers, ex-smokers, and non-smoker COPD individuals compared to healthy controls using plasma samples. Interestingly, we observed metabolic alterations in the smokers and ex-smokers subgroups, except for the non-smokers, underscoring the effect of CS in the development of COPD.

The notable observation of this study is the dysregulation of fatty acids, sugar alcohols, vitamins, fatty acid conjugates, amino acids, and volatile organic compounds (VOCs). Further pathway enrichment analysis reveals perturbation in the biosynthesis of unsaturated fatty acids, GPI anchor metabolism, pyruvate metabolism, fatty acid biosynthesis, and butanoate metabolism. These pathways have been reported to be associated with COPD and lung disease in many studies, such as pyruvate metabolism, which is central to cellular bioenergetics, emphasizing the crisis of energy metabolism in COPD **[15]**. Dysregulation of GPI anchor proteins in alveolar macrophages is known to exacerbate the emphysematous and impaired immune signalling condition in COPD; here we report this for the first time in plasma samples **[16,17]**. Further disruption in fatty acid biosynthesis is well known and reported in the plasma of COPD individuals **[10]**. The fatty acids play a key role in energy production, structural integrity, and signalling of pathways, its alteration likely results in membrane remodelling and immune responses. Butanoate metabolites are derived from gut microbiota, and it exhibits a wide variety of activities, including energy homeostasis, lipid metabolism, and inflammation **[18]**. Interestingly, it has been reported in COPD **[19,20]**.

Pairwise comparison of smokers vs. controls reveals upregulation of α-tocopherol (vitamin E), tetradecanoic acid, tetradecanal, 2,4-di-tert-butylphenol, and 13-docosenamide. The downregulated metabolites were 1-dodecene, chloroxylenol, acetophenone, and docosane. In the ex-smokers subgroup, the upregulated metabolites were lactic acid, 2-hydroxybutyric acid, tetradecanal, 2,4-di-tert-butylphenol, myo-inositol, palmitoleic acid, and ethylene glycol. In contrast, the downregulated compounds were hexadecanoic acid (palmitic acid), dodecanoic acid, scyllo inositol, tetradecanoic acid (myristic acid), octadecanoic acid (stearic acid), 1-dodecene, chloroxylenol, cyclic octatomic sulfur, decane-3-methyl, acetic acid ethenyl ester, and nonane 2,6-dimethyl.

These compounds play roles in fatty acid biosynthesis, antioxidant activity, neuroprotection, and lipid metabolism. For example, α-tocopherol, also known as vitamin E, is a fat-soluble antioxidant that scavenges reactive oxygen species (ROS) and protects the membrane fatty acids from oxidative damage. Increased blood α-tocopherol levels have been correlated with increased oxidative stress **[8]**, which is consistent with our observation. Tetradecanal, a fatty aldehyde, has previously been reported to be upregulated in hepatocellular carcinoma tissue **[21]**. Fatty aldehyde can be produced as an intermediate during the conversion of fatty alcohol to fatty acid by the enzyme fatty aldehyde dehydrogenase or as a result of oxidative damage **[22]**. We observed an increased level of tetradecanal in both the COPD subgroups, implying probable oxidative stress. 2,4-di-tert-butylphenol, an antioxidant and anti-inflammatory molecule **[23,24]**, is known for mitigating oxidative stress; however, it also acts as an endocrine disruptor and results in lipid accumulation **[25]**. Though very limited information is available regarding its role in disease biology and as a biomarker, we observed its increased level in smokers and ex-smokers. 13-docosenamide, a fatty acid amide, has been previously linked with myocardial ischemia and Parkinson’s disease **[26,27]** and is found to be increased in the smoker subgroup. A study by Xu et al. demonstrated the role of 13-docosenamide in cognitive impairment by increasing CNR1 protein, activating phosphorylation of the AKT/mTOR pathway, and promoting oligodendrocyte precursor cell differentiation **[28]**. In addition to this, we observed reduced levels of acetophenone in COPD smokers. Altered levels of acetophenone have been reported previously in COPD and pleural mesothelioma **[9,29,30]**. 1-Dodecene, an unsaturated hydrocarbon which is a product of gut microbiota observed to be reduced in smokers and ex-smokers, and has been previously reported to be downregulated in gastric cancer patients **[31].** Chloroxylenol, which is a known antibiotic and is part of human exposomes observed to be reduced in COPD; there is limited evidence of its role in disease biology in humans; however, studies on colorectal cell lines have demonstrated its anticancerous role **[32]**. Docosane, a long-chain saturated hydrocarbon observed to be downregulated in smokers plasma, indicates elevated oxidative damage associated with CS exposure **[33]**. A study conducted by Yildirim et al, on a COPD mouse model observed the reduced level of docosane in active smokers compared to passive smokers **[34]**. The perturbed level of saturated fatty acids, such as palmitic acid and 2-hydroxybutyric acid, has been associated with various metabolic disorders, such as cardiovascular disorders and lung disorders **[35,36,37,38,39]**. Additionally, it has been previously reported to be decreased in the plasma of COPD patients, consistent with our observation **[40]**. Palmitic acid is proinflammatory and binds to CD36 receptors, stimulating airway cells to produce more immune cells and interleukins through the accumulation of ROS inside the cells **[37]**. This increased inflammatory cascade will further damage the respiratory cells and alveoli. Lactic acid is a metabolic byproduct of glycolysis in an anaerobic environment; increased plasma lactic acid levels reflect muscle fatigue and hypoxia in ex-smokers with COPD **[41,42]**. Scylloinositol, a known sugar alcohol for its neuroprotective role **[43],** has been previously reported to be downregulated in a study conducted by Mandal et al. in the saliva of ex-smoker COPD **[9]**. Tetradecanoic acid (myristic acid) is a saturated fatty acid that is involved in lipid metabolism and protein signalling. We have observed an increase in its level in the smoker subgroup and reduce level in the ex-smoker subgroup. Previously, tetradecanoic acid was reported to be decreased in mild COPD cases, while it was increased in severe COPD patients, suggesting that the COPD individuals who are current smokers have a severe disease condition **[44]**. Another study conducted by Danesh et al. has shown a decrease in tetradecanoic acid in the plasma of COPD patients **[40]**. Further, we observed a reduction in octadecanoic acid (stearic acid) and dodecanoic acid. A decrease in the level of stearic acid has been reported in the plasma of COPD **[40]**. Dodecanoic acid, also known as lauric acid, is a medium-chain saturated fatty acid derived from nutritional diets and triglyceride metabolism and is associated with alzheimer’s disease **[45]**. It is known that CS leads to disruption in glucose and energy metabolism; therefore, to preserve the cellular energy balance, these saturated fatty acids get oxidised to generate energy by the process of fatty acid oxidation, ultimately reducing their level in blood **[46]**. Further reduction in alkane and VOCs in the COPD patients suggests the elevated oxidative stress-driven fragmentation of large-chain hydrocarbons **[33]**.

To further investigate the biological relevance of the altered plasma metabolites, a gene-metabolite interaction network was constructed by integrating the significantly dysregulated metabolites with those genes that were associated with COPD. Several hub genes identified in the network have previously been implicated in the regulation of inflammation, oxidative stress, and mitochondrial dysfunction, all of which are hallmarks of COPD. For example, the FASN (Fatty acid synthase) gene is known to regulate CS-induced oxidative stress, apoptosis, and mitochondrial damage by modulating the composition of the surfactant phospholipidome **[47,48]**. AGXT2 (alanine-glyoxylate aminotransferase 2) gene regulates the L-arginine/nitric oxide (NO) pathway, amino acid metabolism, and oxidative stress in COPD by cleaving ADMA (asymmetric dimethylarginine) and SDMA (symmetric dimethylarginine) **[49,50]**. LGALS13 gene encodes galectin 13, which is related to airway eosinophilic inflammation in COPD **[51,52]**. The interactions between these COPD-associated genes and the altered plasma metabolites suggest that metabolic dysregulation is closely intertwined with the molecular mechanisms driving chronic airway inflammation and tissue remodeling. Although these interactions do not establish direct causal relationships, they provide biologically plausible links between metabolic alterations and established COPD-related pathways.

Overall, this study reveals the perturbation of key metabolic pathways in COPD patients, including biosynthesis of unsaturated fatty acids, GPI anchor metabolism, pyruvate metabolism, fatty acid biosynthesis, and butanoate metabolism. The alteration in these pathways reflects the cumulative implications of CS and disease pathogenesis in smokers and ex-smokers with COPD. Although we have not observed any significant change in non-smoker COPD, this observation should not be interpreted as no metabolic alterations. Further validation in a large and independent cohort is required to establish these findings.

## 5. Conclusion

The present study explores the potential of plasma metabolomics in characterizing metabolic alterations in COPD associated with CS. Our study reveals a disruption in the metabolic profile of smokers and ex-smokers with COPD, implying CS associated disease pathophysiology. Additionally, through ROC analysis, we screened five potential candidate biomarkers, such as tetradecanoic acid, 2,4-di-tert-butylphenol, chloroxylenol, tetradecanal, and 1-dodecene. However, these potential candidate biomarkers require validation in larger, independent cohorts. Overall, this study provides novel insights into the metabolic alterations associated with COPD across different smoking statuses and contributes to the growing understanding of the molecular mechanisms underlying COPD pathogenesis.

## 6. Ethics Declaration

This study was approved by the ethics committee of IISER Kolkata and AIIMS Kalyani (India) with reference number: IEC/AIIMS/Kalyani/certificate/2024/415

## 7. Data availability statement

The data used in this study are available on request from the corresponding author; the remaining data have been provided in the supplementary file.

## 9. CRediT authorship contribution statement

**RS:** Conceptualization, data curation, formal analysis, investigation, methodology, software, visualization, writing-original draft, review and editing. **SG:** Resources, supervision, review and editing. **AKM:** Conceptualization, supervision, resources, writing-review and editing.

## 10. Acknowledgements

The author acknowledges the Council of Scientific and Industrial Research (CSIR), Government of India, for providing the fellowship. We would like to acknowledge all the participants who provided samples for this study, and Namrata Yadav (AIIMS-Kalyani) for helping in sample collection. I would like to acknowledge Neha Yadav (IISER-Kolkata) for her help with the software and valuable guidance during the study. We acknowledge the Nano Mission, Department of Science and Technology, for funding a mass spectrometry facility sanctioned under the project SR/NM/NS-1068/2015. We acknowledge the common instrument facility at IISER Kolkata. We would also like to acknowledge the Department of Earth Science (DES) at IISER Kolkata for providing access to the GC-MS instrument.

## References

1. Heffler, E., Crimi, C., Mancuso, S., Campisi, R., Puggioni, F., Brussino, L., & Crimi, N. (2018). Misdiagnosis of asthma and COPD and underuse of spirometry in primary care unselected patients. Respiratory medicine, 142, 48–52. DOI 10.1016/j.rmed.2018.07.015.

2. Vaz Fragoso, C. A., McAvay, G., Van Ness, P. H., Casaburi, R., Jensen, R. L., MacIntyre, N., … & Concato, J. (2016). Phenotype of spirometric impairment in an aging population. American journal of respiratory and critical care medicine, 193(7), 727–735. DOI 10.1164/rccm.201508-1603OC.

3. Halper-Stromberg, E., Gillenwater, L., Cruickshank-Quinn, C., O’Neal, W. K., Reisdorph, N., Petrache, I., Zhuang, Y., Labaki, W. W., Curtis, J. L., Wells, J., Rennard, S., Pratte, K. A., Woodruff, P., Stringer, K. A., Kechris, K., & Bowler, R. P. (2019). Bronchoalveolar Lavage Fluid from COPD Patients Reveals More Compounds Associated with Disease than Matched Plasma. Metabolites, 9(8), 157. DOI 10.3390/metabo9080157

4. Banerjee, S., Bhattacharyya, P., Mitra, S., Kundu, S., Panda, S., & Chatterjee, I. B. (2017). Anti-p-benzoquinone antibody level as a prospective biomarker to identify smokers at risk for COPD. International Journal of Chronic Obstructive Pulmonary Disease, 12, 1847. DOI 10.2147/COPD.S134455.

5. Yadav, N., Singh, R., Mondal, S. K., & Mandal, A. K. (2026). Molecular Insights of p-Benzoquinone-Induced Red Blood Cell Dysfunction: Probable Implications to Cigarette Smoke-Associated Pathologies. Free Radical Research, 60(1) , 67–90. DOI 10.1080/10715762.2026.2620638

6. Yadav, N., PM, J., Mondal, S. K., & Mandal, A. K. (2025). Differential Reactivity of Airborne Quinones on Human Red Blood Cells: Insights into Their Biochemical and Morphological Alterations. Chemical Research in Toxicology, 38(11), 1984–2001. DOI 10.1021/acs.chemrestox.5c00304

7. Kabir, E., Raza, N., Kumar, V., Singh, J., Tsang, Y. F., Lim, D. K., … & Kim, K. H. (2019). Recent advances in nanomaterial-based human breath analytical technology for clinical diagnosis and the way forward. Chem, 5(12), 3020–3057. DOI 10.1016/j.chempr.2019.08.004

8. Miranda, C. T. D. O. F., Duarte, V. H. R., Cruz, M. S. D. M., Duarte, M. K. R. N., de Araújo, J. N. G., Santos, A. M. Q. S. D., … & Silbiger, V. N. (2018). Association of Serum Alpha-Tocopherol and Retinol with the Extent of Coronary Lesions in Coronary Artery Disease. Journal of nutrition and metabolism, 2018(1), 6104169. DOI 10.1155/2018/6104169

9. Singh, R., Ghosh, S., Yadav, N., & Mandal, A. K. (2026). Saliva Metabolomics Reveals Distinct Metabolic Signatures in Patients with Chronic Obstructive Pulmonary Disease: A GC-MS-based approach. bioRxiv, (Preprint). DOI 10.64898/2026.04.10.717654

10. Casadevall, C., Agranovich, B., Enríquez-Rodríguez, C. J., Faner, R., Pascual-Guardia, S., Castro-Acosta, A., … & Gea, J. (2025). Metabolomic plasma profile of chronic obstructive pulmonary disease patients. International journal of molecular sciences, 26(10), 4526. DOI 10.3390/ijms26104526.

11. Zhou, J., Li, Q., Liu, C., Pang, R., & Yin, Y. (2020). Plasma metabolomics and lipidomics reveal perturbed metabolites in different disease stages of chronic obstructive pulmonary disease. International Journal of Chronic Obstructive Pulmonary Disease, 553–565. DOI 10.2147/COPD.S229505.

12. Godbole, S., & Bowler, R. P. (2022). Metabolome Features of COPD: A Scoping Review. Metabolites, 12(7), 621. DOI 10.3390/metabo12070621

13. Lv, J., Pan, C., Cai, Y., Han, X., Wang, C., Ma, J., … & Chen, Y. (2024). Plasma metabolomics reveals the shared and distinct metabolic disturbances associated with cardiovascular events in coronary artery disease. Nature Communications, 15(1), 5729. DOI 10.1038/s41467-024-50125-2

14. Liu, Z., Wang, L., Gao, S., Xue, Q., Tan, F., & Gao, Y. (2025). Plasma metabolites as biomarkers for screening and differential diagnosis of lung cancer. Journal of Cardiothoracic Surgery, 20(1), 456. DOI 10.1186/s13019-025-03719-w

15. Zeng, S., Zhang, Y., Li, S., Li, Z., Li, P., Xie, J., Zhang, J., Xie, L., & Yang, Y. (2025). From metabolic alterations to chronic inflammation: mechanisms and immunoregulation of metabolic reprogramming in COPD. Frontiers in Immunology, 16. DOI:10.3389/fimmu.2025.1698832

16. Ponessa, J. J., Peng, J., Schroeter, M. N., Sawhney, A., Chavan, K., Rico, J. I., Espinosa, V., Chen, F., Shubin, N. J., Wong Fok Lung, T., Lemenze, A., Rivera, A., Gause, W. C., & Siracusa, M. C. (2026). Alveolar macrophages inhibit emphysematous pathology via expression of carbonic anhydrase 4. Cell Reports, 45(5), 117315. DOI 10.1016/j.celrep.2026.117315

17. Wen, H., Zhang, R., Zhong, B., Liu, H., & Liu, C. (2025). Cross-Trait Genome-Wide Association Study Identifies Shared Genetic Risk Loci Between COPD and Five Autoimmune Diseases. International Journal of Chronic Obstructive Pulmonary Disease, Volume 20, 3019–3034. DOI 10.2147/COPD.S533401

18. Amiri, P., Hosseini, S. A., Ghaffari, S., Tutunchi, H., Ghaffari, S., Mosharkesh, E., … & Roshanravan, N. (2022). Role of butyrate, a gut microbiota derived metabolite, in cardiovascular diseases: a comprehensive narrative review. Frontiers in Pharmacology, 12, 837509. DOI 10.3389/fphar.2021.837509

19. Feng, Y., Xie, M., Liu, Q., Weng, J., Wei, L., Chung, K. F., Adcock, I. M., Chang, Q., Li, M., Huang, Y., Zhang, H., & Li, F. (2023). Changes in targeted metabolomics in lung tissue of chronic obstructive pulmonary disease. Journal of Thoracic Disease, 15(5), 2544–2558. DOI 10.21037/jtd-22-1731

20. Gea, J., Enríquez-Rodríguez, C. J., Agranovich, B., & Pascual-Guardia, S. (2023). Update on metabolomic findings in COPD patients. ERJ Open Research, 9(5), 00180–02023. DOI 10.1183/23120541.00180-2023

21. Liu, S. Y., Zhang, R. L., Kang, H., Fan, Z. J., & Du, Z. (2013). Human liver tissue metabolic profiling research on hepatitis B virus-related hepatocellular carcinoma. World journal of gastroenterology: WJG, 19(22), 3423. DOI 10.3748/wjg.v19.i22.3423

22. Lloyd, M. D., Boardman, K. D., Smith, A., Van Den Brink, D. M., Wanders, R. J., & Threadgill, M. D. (2007). Characterisation of recombinant human fatty aldehyde dehydrogenase: implications for Sjögren-Larsson syndrome. Journal of Enzyme Inhibition and Medicinal Chemistry, 22(5), 584–590. DOI 10.1080/14756360701425360

23. Vahdati, S. N., Lashkari, A., Navasatli, S. A., Ardestani, S. K., & Safavi, M. (2022). Butylated hydroxyl-toluene, 2, 4-Di-tert-butylphenol, and phytol of Chlorella sp. protect the PC12 cell line against H2O2-induced neurotoxicity. Biomedicine & Pharmacotherapy, 145, 112415. DOI 10.1016/j.biopha.2021.112415

24. Eleazu, C. O., Obeten, U. N., Ozor, G., Njemanze, C. C., Eleazu, K. C., Egedigwe-Ekeleme, A. C., … & Kanu, S. (2022). Tert-butylhydroquinone abrogates fructose-induced insulin resistance in rats via mitigation of oxidant stress, NFkB-mediated inflammation in the liver but not the skeletal muscle of high fructose drinking rats. Journal of Food Biochemistry, 46(12), e14473. DOI 10.1111/jfbc.1447

25. Ren, X. M., Chang, R. C., Huang, Y., Amorim Amato, A., Carivenc, C., Grimaldi, M., … & Blumberg, B. (2023). 2, 4-Di-tert-butylphenol induces adipogenesis in human mesenchymal stem cells by activating retinoid X receptors. Endocrinology, 164(4), bqad021. DOI 10.1210/endocr/bqad021

26. Gątarek, P., Sekulska-Nalewajko, J., Bobrowska-Korczaka, B., Pawełczyk, M., Jastrzębski, K., Gąbiński, A., & Kałuńa-Czaplińska, J. (2022). Plasma metabolic disturbances in Parkinson’s disease patients. Biomedicines, 10(12), 3005. DOI 10.3390/biomedicines10123005

27. Han, H. J., Gwon, M. R., Lee, J. H., Park, J. S., Lee, H. W., Yoon, Y. R., … & Seong, S. J. (2026). Plasma metabolomic profiling reveals lipid biomarkers for early detection of exercise-induced myocardial ischemia. Medicine, 105(12), e47995. DOI 10.1097/MD.0000000000047995

28. Xu, Y., Tan, Y., Zhang, Z., Chen, D., Zhou, C., Sun, L., … & Xu, Y. (2025). 13- Docosenamide Enhances Oligodendrocyte Precursor Cell Differentiation via USP33-Mediated Deubiquitination of CNR1 in Chronic Cerebral Hypoperfusion: Y. Xu et al.: 13-Docosenamide Enhances Oligodendrocyte Precursor Cell Differentiation. Neuroscience Bulletin, 41(11), 1939–1956. DOI 10.1007/s12264-025-01461-w

29. de Gennaro, G., Dragonieri, S., Longobardi, F., Musti, M., Stallone, G., Trizio, L., & Tutino, M. (2010). Chemical characterization of exhaled breath to differentiate between patients with malignant plueral mesothelioma from subjects with similar professional asbestos exposure. Analytical 398(7),3043–3050. DOI 10.1007/s00216-010-4238-y.

30. Di Gilio, A., Catino, A., Lombardi, A., Palmisani, J., Facchini, L., Mongelli, T., … & Tangaro, S. (2020). Breath analysis for early detection of malignant pleural mesothelioma: volatile organic compounds (VOCs) determination and possible biochemical pathways. Cancers,12(5), 1262. DOI 10.3390/cancers12051262.

31. Bhandari, M. P., Polaka, I., Vangravs, R., Mezmale, L., Veliks, V., Kirshners, A., … & Leja, M. (2023). Volatile markers for cancer in exhaled breath—could they be the signature of the gut microbiota?. Molecules, 28(8), 3488. DOI 10.3390/molecules28083488

32. Sun, Q., Liu, B., Lan, Q., Su, Z., Fu, Q., Wang, L., … & Lu, D. (2023). Antimicrobial agent chloroxylenol targets â-catenin-mediated Wnt signaling and exerts anticancer activity in colorectal cancer. International Journal of Oncology, 63(5), 121. DOI 10.3892/ijo.2023.5569

33. Van Berkel, J. J. B. N., Dallinga, J. W., Möller, G. M., Godschalk, R. W. L., Moonen, E. J., Wouters, E. F. M., & Van Schooten, F. J. (2010). A profile of volatile organic compounds in breath discriminates COPD patients from controls. Respiratory medicine, 104(4),557–563. DOI 10.1016/j.rmed.2009.10.018

34. John, G., Kohse, K., Orasche, J., Reda, A., Schnelle-Kreis, J., Zimmermann, R., … & Yildirim, A. Ö. (2014). The composition of cigarette smoke determines inflammatory cell recruitment to the lung in COPD mouse models. Clinical science, 126(3), 207–221. DOI 10.1042/CS20130117

35. Li, Z., Lei, H., Jiang, H., Fan, Y., Shi, J., Li, C., … & Ma, L. (2022). Saturated fatty acid biomarkers and risk of cardiometabolic diseases: A meta-analysis of prospective studies. Frontiers in nutrition, 9, 963471. DOI 10.3389/fnut.2022.963471

36. Saraswathi, V., Kumar, N., Ai, W., Gopal, T., Bhatt, S., Harris, E. N., … & Desouza, C. V. (2022). Myristic acid supplementation aggravates high fat diet-induced adipose inflammation and systemic insulin resistance in mice. Biomolecules 2022; 12: 739. DOI 10.3390/biom12060739

37. Wu, Y., Ma, J., Shan, C., Li, W., Chen, Q., Miao, X., … & Ni, Z. (2026). Palmitic acid aggravates airway inflammation in asthma through induction of chemokines expression and metabolomic changes in epithelial cells. Allergology International. DOI 10.1016/j.alit.2026.01.004

38. Qin, F., Li, J., Mao, T., Feng, S., Li, J., & Lai, M. (2023). 2 hydroxybutyric acid-producing bacteria in gut microbiome and fusobacterium nucleatum regulates 2 hydroxybutyric acid level in vivo. Metabolites, 13(3), 451. DOI 10.3390/metabo13030451

39. Shima, N., Miki, A., Kamata, T., Katagi, M., & Tsuchihashi, H. (2005). Urinary endogenous concentrations of GHB and its isomers in healthy humans and diabetics. Forensic science international, 149(2-3), 171–179. DOI 10.1016/j.forsciint.2004.05.017

40. Yazdani, R., Fallah, H., Yazdani, S., Shahouzehi, B., & Danesh, B. (2025). Effect of plasma free fatty acids on lung function in male COPD patients. Scientific Reports, 15(1), 3377. DOI 10.1038/s41598-025-86628-1

41. Wang, H., Cho, P. S., Kouritas, V., & Feng, L. (2025). Three key factors predicting the severity of exacerbations of chronic obstructive pulmonary disease: T lymphocytes, lactate, and prealbumin. Journal of Thoracic Disease, 17(4), 2386–2393. DOI 10.21037/jtd-2025-416

42. Wu LW, Kao TW, Lin CM, Yang HF, Sun YS, Liaw FY, et al. Examining the association between serum lactic dehydrogenase and all-cause mortality in patients with metabolic syndrome: A retrospective observational study. BMJ Open. 2016;6:e011186. DOI 10.1136/bmjopen-2016-011186.

43. Ma, K., Thomason, L. A., & McLaurin, J. (2012). Scyllo-inositol, preclinical, and clinical data for Alzheimer’s disease. Advances in pharmacology, 64, 177–212. DOI 10.1016/B978-0-12-394816-8.00006-4.

44. Novgorodtseva, T. P., Vitkina, T. I., Knyshova, V. V., Antonyuk, M. V., Bocharova, N. V., & Kytikova, O. Y. (2022). Associations of fatty acid composition in leukocyte membranes with systemic inflammation in chronic obstructive pulmonary disease progression. Russian Open Medical Journal, 11(4), 401. DOI 10.15275/rusomj.2022.0401

45. Jasbi, P., Shi, X., Chu, P., Elliott, N., Hudson, H., Jones, D., … & Gu, H. (2021). Metabolic profiling of neocortical tissue discriminates Alzheimer’s disease from mild cognitive impairment, high pathology controls, and normal controls. Journal of proteome research, 20(9), 4303–4317. DOI 10.1021/acs.jproteome.1c00290

46. Liang, Q., Wang, Y., & Li, Z. (2025). Lipid metabolism reprogramming in chronic obstructive pulmonary disease. Molecular Medicine, 31(1), 129. DOI 10.1186/s10020-025-01191-9

47. Yang, K., Zhu, G., Peng, T., Cheng, Y., & Guo, X. (2025). FASN regulates CSE-induced apoptosis, oxidative stress and mitochondrial damage in type 2 alveolar epithelial cells by regulating NRF2 expression and nuclear translocation. Redox Report, 30(1), 2550412. DOI 10.1080/13510002.2025.2550412

48. Fan, L. C., McConn, K., Plataki, M., Kenny, S., Williams, N. C., Kim, K., … & Cloonan, S. M. (2023). Alveolar type II epithelial cell FASN maintains lipid homeostasis in experimental COPD. JCI insight, 8(16), e163403. DOI 10.1172/jci.insight.163403

49. Hannemann, J., & Böger, R. (2022). Dysregulation of the nitric oxide/Dimethylarginine pathway in hypoxic pulmonary vasoconstriction—molecular mechanisms and clinical significance. Frontiers in Medicine, 9, 835481. DOI 10.3389/fmed.2022.835481

50. Belskikh, E., Marsyanova, Y., Melnikov, D., Uryasev, O., & Zvyagina, V. (2025). Endothelial and Mitochondrial Dysfunction in COPD Pathophysiology: Focus on Homocysteine–L-Carnitine Interplay. Biocell, 49(11). DOI 10.32604/biocell.2025.069272

51. Yi, L., Feng, Y., Chen, D., Jin, Y., & Zhang, S. (2023). Association between galectin-13 expression and eosinophilic airway inflammation in chronic obstructive pulmonary disease. COPD: Journal of Chronic Obstructive Pulmonary Disease, 20(1), 101–108. DOI 10.1080/15412555.2022.2162377

52. Zhao, X., Han, B., Tang, W., Ji, S., Wang, L., Huang, J., … & Li, J. (2024). Association between serum galectin-3 and chronic obstructive pulmonary disease: A meta-analysis. Biomolecules and Biomedicine, 24(6), 1491. DOI 10.17305/bb.2024.10527

