## Supplemental data for "Plasma Metabolomic Profiling of COPD Patients Stratified by Smoking Status: A GC-MS-Based Approach"

### Table of contents:

#### Supplemental Tables

**Supplementary Table S1:** List of significantly altered metabolites between COPD subgroups and healthy controls.

**Supplementary Table S2:** List of significantly altered metabolites between COPD smokers and healthy controls.

**Supplementary Table S3:** List of significantly altered metabolites between COPD ex-smokers and healthy controls

#### Supplemental Figures

**Supplementary Figure S1:** Heatmap profile of differential metabolites in COPD smokers vs healthy controls.

**Supplementary Figure S2:** Heatmap profile of differential metabolites in COPD ex-smokers vs healthy controls.

**Supplementary Figure S3:** ROC curve plot metabolites in smokers vs control analysis

**Supplementary Figure S3:** ROC curve plot metabolites in ex-smokers vs control analysis

**Table S1:** List of significantly altered metabolites between COPD subgroups and Healthy controls with VIP >1, p-value <0.05, FDR <0.05, and AUC>0.8.

| S.No | Metabolites | HMDB ID | Pubchem ID | p-value$_{Adj}$ | VIP | AUC |
|---|---|---|---|---|---|---|
| 1 | 2-Benzyl-1,3-oxazol-2-ine | | | 2.41E-05 | 2.6148 | 0.983333 |
| 2 | Chloroxylenol | HMDB0246387 | 2723 | 2.53E-05 | 1.7588 | 0.95 |
| 3 | 5-Iodopentan-2-one | | 538269 | 2.56E-05 | 1.7748 | 0.945833 |
| 4 | Hexathiane | | | 4.49E-05 | 1.6328 | 0.920833 |
| 5 | 2,6-Difluorobenzoic acid, 2-naphthyl ester | | | 5.37E-05 | 2.0874 | 0.9125 |
| 6 | Dodecanoic acid | HMDB0000638 | 3893 | 5.73E-05 | 1.1434 | 0.883333 |
| 7 | 1-Dodecene | HMDB0059874 | 8183 | 7.85E-05 | 1.9038 | 0.916667 |
| 8 | Tetradecanal, O-methyloxime | HMDB0034283 | 31291 | 0.000292 | 1.447 | 0.870833 |
| 9 | Piperazine-2,5-dione, 1,4-dimethyl-3,3'-bis- | | 78761 | 0.000317 | 1.2383 | 0.820833 |
| 10 | 2-Amino-2-oxoethyl acetate | | | 0.001042 | 1.3129 | 0.883333 |

| | | | | | | |
|---|---|---|---|---|---|---|
| 11 | Acetone | HMDB0001 659 | 180 | 0.0014 6 | 1.37 99 | 0.8083 33 |
| 12 | Nonane, 4,5-dimethyl- | | 86541 | 0.0016 4 | 1.43 98 | 0.8291 67 |
| 13 | Cyclobutene-3,4-dione, 1-dimethylamino-2-hydroxy- | | | 0.0016 8 | 1.69 41 | 0.8833 33 |
| 14 | Methyl hydrogen disulfide | | 52205 9 | 0.0020 21 | 1.70 88 | 0.9208 33 |
| 15 | Decane, 3-methyl- | HMDB0037 267 | 92239 | 0.0048 8 | 1.46 82 | 0.8541 67 |
| 16 | Undecane, 4,7-dimethyl | | 51938 9 | 0.0107 64 | 1.30 47 | 0.8333 33 |
| 17 | Phenol, 2,4-bis(1,1-dimethylethyl)-, phosphite (3:1) | | 26004 2 | 0.0110 28 | 1.27 73 | 0.8458 33 |
| 18 | 3-Trifluoromethylbenzylamine, N,N-dinonyl | | | 0.0122 33 | 1.55 86 | 0.8833 33 |

**Table S2:** List of significantly altered metabolites between COPD smokers and Healthy controls with VIP >1, p-value <0.05, FDR <0.05, AUC>0.8, and $Log_2FC \geq \pm 1$.

| S.No | Metabolites | HMDB ID | Pubchem ID | p-value$_{Adj}$ | VIP | Log2 FC | AUC |
|---|---|---|---|---|---|---|---|
| 1 | Benzene, 1,1'-[oxybis(methylene)]bis- | HMDB0032078 | 31232 | 9.18E-05 | 2.10742 | 4.31 | 1 |
| 2 | Tetradecanoic acid | HMDB0000806 | | 9.18E-05 | 1.052149 | 1.48 | 1 |
| 3 | 2,6-Difluorobenzoic acid, 2-naphthyl ester | | | 9.18E-05 | 1.911113 | 3.60 | 1 |
| 4 | dl-.alpha.-Tocopherol | HMDB0001893 | | 9.18E-05 | 2.259499 | 5.20 | 1 |
| 5 | 2-Benzyl-1,3-oxazol-2-ine | | | 9.18E-05 | 2.528326 | -11.1 | 1 |
| 6 | 13-Docosenamide, (Z)- | HMDB0244507 | | 0.000734 | 1.296668 | 2.4 | 0.96 |
| 7 | Chloroxylenol | HMDB0246387 | | 0.000734 | 1.830661 | -3.5 | 0.96 |
| 8 | 1-Dodecene | HMDB0059874 | | 0.001073 | 1.951832 | -2.91 | 0.95 |
| 9 | Cyclobutene-3,4-dione, 1- | | | 0.001573 | 1.990488 | -3.03 | 0.95 |

| | | | | | | | |
|---|---|---|---|---|---|---|---|
| | dimethylamino -2-hydroxy- | | | | | | |
| 10 | 5-Iodopentan-2-one | | 538269 | 0.002203 | 1.886 055 | - 4.44 | 0.94 |
| 11 | Benzene, (isocyanometh y) | | 76639 | 0.002236 | 1.027 898 | 1.77 | 0.93 |
| 12 | 2,2'-Bi-1,3-dioxolane | | | 0.002236 | 1.670 724 | 3.22 | 0.93 |
| 13 | Cholesta-3,5-diene | | 92835 | 0.002236 | 1.559 469 | 3.35 | 0.93 |
| 14 | Methyl hydrogen disulfide | | | 0.002236 | 1.845 679 | - 5.35 | 0.93 |
| 15 | Pentane, 3,3-dimethyl- | | 11229 | 0.002236 | 2.455 575 | - 1.44 | 0.93 |
| 16 | Benzyl Benzoate | HMDB 001481 4 | | 0.002236 | 1.940 108 | - 2.18 | 0.93 |
| 17 | 3-Trifluoromethy lbenzylamine, N,N-dinonyl | | | 0.003065 | 1.943 664 | - 2.44 | 0.92 |
| 18 | 2,4-Di-tert-butylphenoxytr imethylsilane | HMDB 001381 6 | 7311 | 0.003994 | 2.015 946 | 4.07 | 0.91 |
| 19 | Acetophenone | HMDB 003391 0 | | 0.007721 | 1.017 686 | - 2.59 | 0.89 |
| 20 | Docosane | HMDB | | 0.012775 | 1.855 | - | 0.88 |

| | | | | | | | |
|---|---|---|---|---|---|---|---|
| | | 0061865 | | | 497 | 1.74 | |
| 21 | Diethyl Phthalate | HMDB0094660 | | 0.019747 | 1.12593 | 1.23 | 0.85 |
| 22 | Hexathiane | | | 0.019747 | 1.865507 | -2.2 | 0.85 |
| 23 | Tetradecanal, O-methyloxime | HMDB0034283 | | 0.047339 | 1.043012 | 2.2 | 0.82 |

**Table S3:** List of significantly altered metabolites between COPD Ex-smoker and Healthy controls having VIP >1, p-value <0.05, FDR <0.05, AUC>0.8, and $Log_2FC \geq \pm 1$.

| **S.No.** | **Metabolites** | **HMDB ID** | **Pubchem ID** | p-value$_{Adj}$ | VIP | Log2FC | AUC |
|---|---|---|---|---|---|---|---|
| 1 | Ethylene glycol, 2TMS derivative | HMDB0037790 | 174 | 0.000188 | 2.181516 | 6.4466 | 1 |
| 2 | Lactic Acid, 2TMS derivative | HMDB0000190 | 107689 | 0.000188 | 1.894273 | 5.6658 | 1 |
| 3 | 2-Hydroxybutyric acid, 2TMS | HMDB0000008 | 440864 | 0.000188 | 2.379046 | 9.9027 | 1 |
| 4 | Tetradecanal, O-methyloxime | HMDB0034283 | 31291 | 0.000188 | 1.645924 | 4.055 | 1 |
| 5 | n-Hexadecanoic acid | HMDB0000220 | 985 | 0.000188 | 2.013923 | -6.4478 | 1 |

| | | | | | | | |
|---|---|---|---|---|---|---|---|
| 6 | Benzene, (isocyanomethyl)- | | 76639 | 0.000188 | 2.410495 | -10.491 | 1 |
| 7 | 2-Benzyl-1,3-oxazol-2-ine | | | 0.000188 | 2.143564 | -12.223 | 1 |
| 8 | 2,4-Di-tert-butylphenoxytrimethylsilane | HMDB0013816 | 7311 | 0.000188 | 2.146074 | 5.1642 | 1 |
| 9 | Hexathiane | | | 0.000188 | 2.41771 | -11.552 | 1 |
| 10 | Dodecanoic acid | HMDB0000638 | 3893 | 0.000188 | 2.313873 | -9.9028 | 1 |
| 11 | Diethyleneglycol adipate | | | 0.000188 | 2.055584 | -8.4699 | 1 |
| 12 | Tetradecanoic acid | HMDB0000806 | 11005 | 0.000188 | 2.237834 | -9.3027 | 1 |
| 13 | 5-Iodopentan-2-one | | | 0.000188 | 1.827984 | -10.746 | 1 |
| 14 | Scyllo-Inositol,6TMS | HMDB0006088 | 892 | 0.000188 | 1.976376 | -10.097 | 1 |
| 15 | 3-(4-Fluorophenyl)-1H-pyrazole-5-carboxylic acid | | | 0.000188 | 2.064686 | -8.6879 | 1 |
| 16 | Octadecanoic acid | HMDB0000827 | 5281 | 0.000188 | 2.340168 | -10.493 | 1 |
| 17 | Chloroxylenol | HMDB0246387 | 2723 | 0.000188 | 1.818893 | -7.4723 | 1 |
| 18 | Hexadecane, 2,6,10,14- | | 12523 | 0.000718 | 1.4084 | 2.9865 | 0.97 |

| | | | | | | | |
|---|---|---|---|---|---|---|---|
| | tetramethyl- | | | | | | |
| 19 | Piperazine-2,5-dione,1,4-dimethyl-3,3'-bis- | | | 0.001204 | 2.387704 | -5.4038 | 0.96 |
| 20 | Edetic Acid, 4TMS derivative | HMDB0015109 | 6049 | 0.001905 | 1.120458 | 3.9596 | 0.95 |
| 21 | Myo-Inositol, 6TMS | HMDB0000211 | | 0.002801 | 1.442605 | 3.562 | 0.94 |
| 22 | 1-Dodecene | HMDB0059874 | 8183 | 0.002801 | 1.658608 | -2.3088 | 0.94 |
| 23 | Silanol, trimethyl-, phosphate (3:1) | HMDB0001429 | | 0.005992 | 1.554806 | 4.9944 | 0.92 |

| | | | | | | | |
|---|---|---|---|---|---|---|---|
| 24 | 9-Hexadecanoic acid, (Z)-, TBDMS derivative | HMDB0003229 | 445638 | 0.005992 | 1.517598 | 4.824 | 0.92 |
| 25 | Ethanol,2-ethoxy | HMDB0031213 | | 0.005992 | 2.001826 | -3.1428 | 0.92 |
| 26 | 1,3-Propanediol, 2TMS derivative | METPA0292 | | 0.007092 | 1.633141 | 5.7235 | 0.91 |
| 27 | 6H-Dibenzo(b,d)pyran-1-ol, 3-hexyl-7,8,9,10-tetrahydro-6,6,9-trimethyl | | | 0.007092 | 1.748537 | 6.3582 | 0.91 |
| 28 | Anthraquinone,1-(o-chlorophenyl)- | | | 0.007092 | 1.157383 | 3.3855 | 0.91 |
| 29 | Cyclic-octaatomic sulfur | HMDB0302183 | | 0.007092 | 1.813233 | -4.5845 | 0.91 |
| 30 | Acetone | HMDB0001659 | 180 | 0.009666 | 1.55532 | -2.4367 | 0.90 |
| 31 | 2-Amino-2-oxoethyl acetate | | | 0.009666 | 1.444645 | -2.4187 | 0.90 |
| 32 | (4-chlorobutoxy) trimethylsilane | | | 0.012476 | 1.481356 | 1.9697 | 0.89 |
| 33 | Methyl hydrogen disulfide | | 522059 | 0.012476 | 1.595343 | -4.9201 | 0.89 |

| | | | | | | | |
|---|---|---|---|---|---|---|---|
| 34 | Acetic acid ethenyl ester | HMDB0031209 | 7904 | 0.021735 | 1.656672 | -4.3355 | 0.87 |
| 35 | Decane, 3-methyl | HMDB0037267 | 92239 | 0.028566 | 1.300615 | -1.347 | 0.86 |
| 36 | Nonane,2,6-dimethyl- | | 28457 | 0.035828 | 1.830836 | -1.4323 | 0.85 |

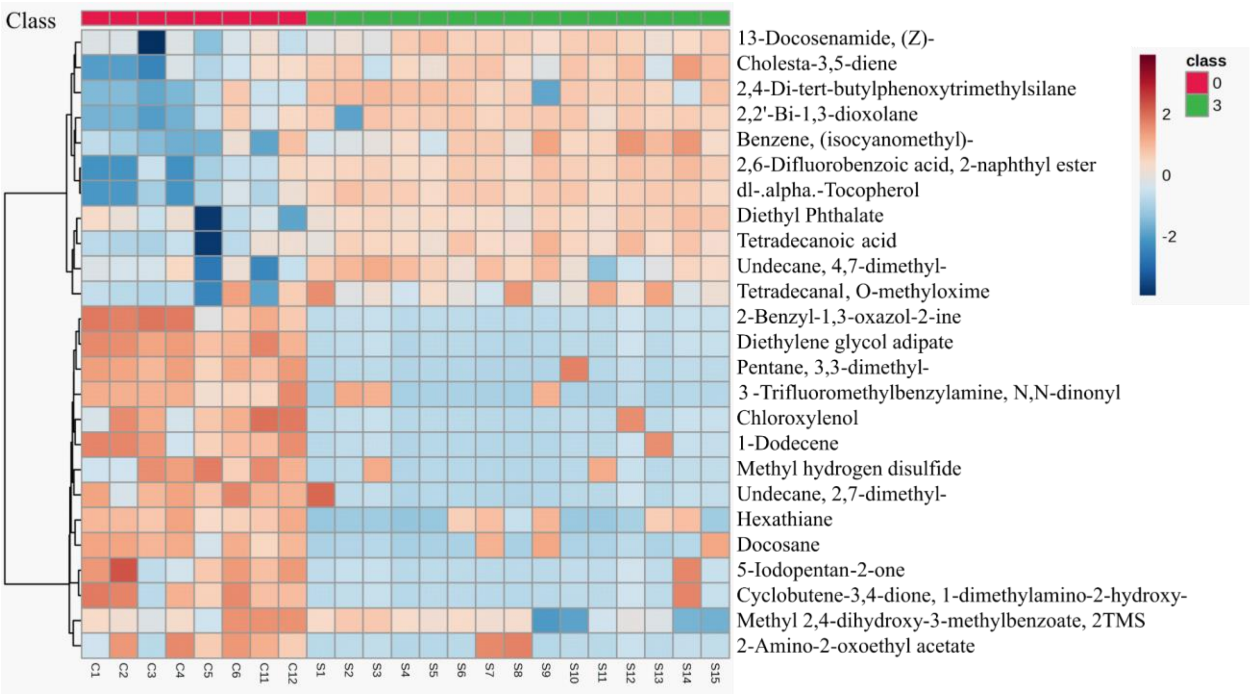


**Figure S1.** Heatmap depicting hierarchical clustering with differentially expressed metabolites of COPD smoker subgroups and healthy controls with p-value < 0.05 (class 0- Healthy control, class 3- Smoker). The red-colored bars show upregulated metabolites, and the blue-colored bars represent downregulated metabolites.

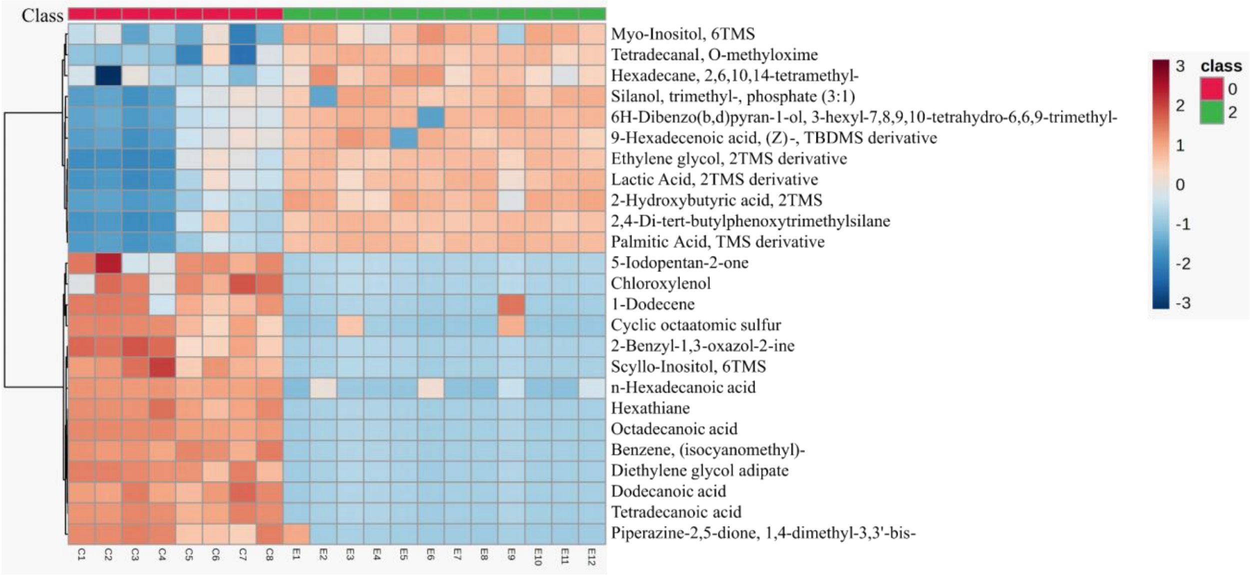


**Figure S2.** Heatmap illustrating hierarchical clustering of differentially expressed metabolites in COPD ex-smokers subgroup and healthy controls with p-value < 0.05 (class 0: Healthy control, class 2: Ex-smoker). The red-colored bars show upregulated metabolites, and the blue-colored bars represent downregulated metabolites.

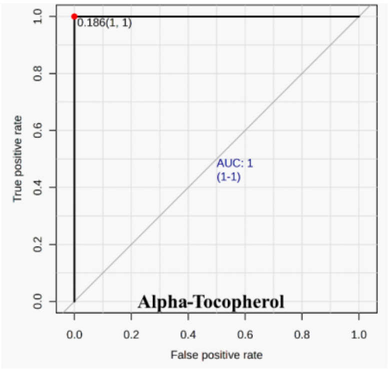


A

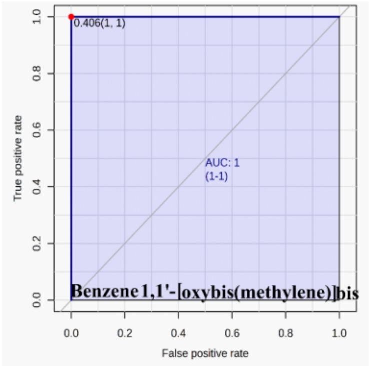


B

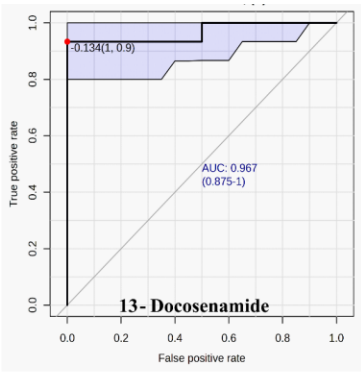


C

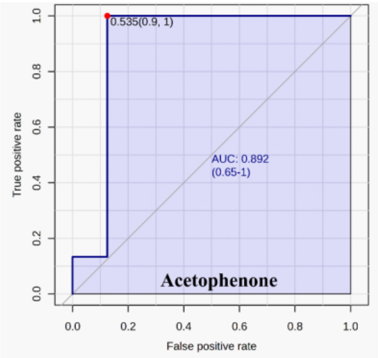


D

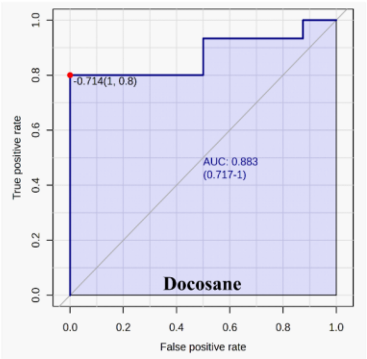


E

**Figure S3:** Receiver operating characteristic curve of top 5 metabolites of smokers COPD subgroup with AUC >0.85.

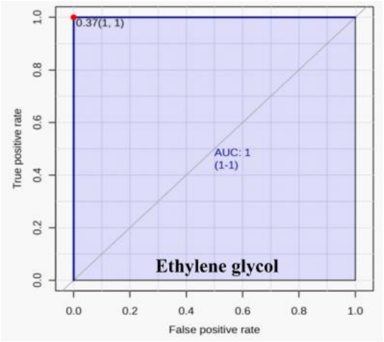


A

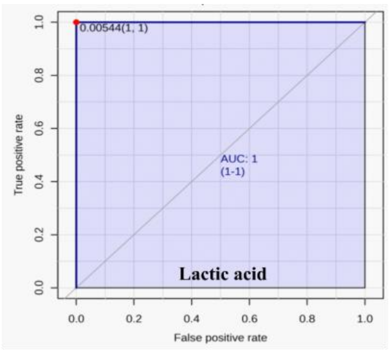


B

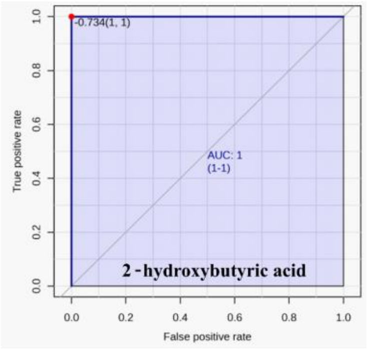


C

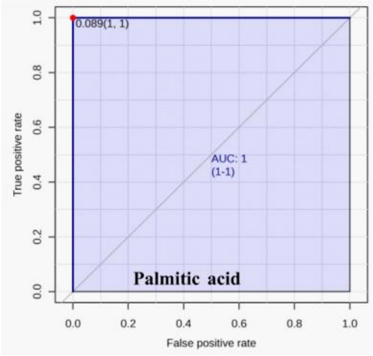


D

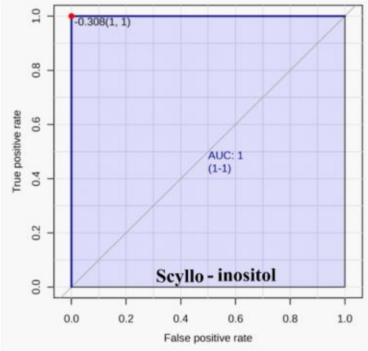


E

**Figure S4:** Receiver operating characteristic curve of top 5 metabolites of ex-smokers COPD subgroup with AUC >0.85.
